# Optimization of an automated system (ZEG) for rapid cellular extraction from live zebrafish

**DOI:** 10.64898/2026.02.19.706735

**Authors:** Nusrat Tazin, Christopher Jordon Lambert, Raheel Samuel, Sabin Nepal, Bruce K Gale

## Abstract

Collecting cells from zebrafish embryos for genotyping is critical to rapid research with these model organisms. The standard collection process is manual, labor-intensive, time-consuming, and requires a skilled person to perform it. To overcome this challenge, researchers are exploring the development of automated genotyping tools for live animals, which would significantly enhance the efficiency and accuracy of genetic screening in zebrafish and other species. The focus of this research was to optimize the Zebrafish Embryo Genotyper (ZEG), an automated system used for the rapid extraction of cellular material from zebrafish embryos. This system rapidly vibrates a roughened chip containing a zebrafish embryo to collect genetic material safely and efficiently. The aim was to improve the efficiency of DNA collection from the chips used with the ZEG by identifying the key factors that contribute to the process. First, the chips were modified to resolve issues associated with loss of sample volume from the chip wells due to evaporation during processing. Second, we experimented with three critical parameters – sample volume in the wells, the vibrational frequency of the system, and the operation time – on the quantity of DNA collected. The performance was evaluated by measuring embryo survival and quantifying the DNA collected. The sensitivity (previously 90%) of the DNA collection and embryo survival (previously 95%) of the were both found to be greater than 95% after optimization. The optimized design parameters (15 µL solution volume, 2.4 V, and a 5-minute run with 5 s alternating on/off) provided a >50% increase in DNA collection compared to the previous designs and parameters. The proposed chip design and operation do not appear to cause any apparent adverse effects on the development or survival of the embryos.

## 1. Introduction

Zebrafish (*Danio rerio*) are considered a leading vertebrate model and are used extensively by the biomedical research community for developmental biology [1] research, disease modeling, and drug discovery [2]. This is largely due to a common genetic and drug toxicity profile similarity with mammals [2][3][4], embryo development outside the mother [4], optical transparency [5], fast development to adulthood [2][5], easy handling, and low-cost maintenance. One of the significant features of the zebrafish model is its amenability to genetic manipulation, which can be improved by incorporating automated research tools. This will reduce reliance on skilled and labor-intensive work, which is prone to result in corrupted outcomes.

The large-scale screening of mutagenesis in zebrafish or determining the transgenic nature of a fish is limited by the labor and time-intensive process, as well as the absence of automated genotyping tools for live animals. Conventionally, genotyping involves raising the transfected zebrafish embryos to adulthood (2-3 months) and then manually clipping their fins by a trained technician. This requires nursery space and maintenance costs, and the throughput of fin clipping and genotyping is limited to around 30 embryos per hour for skilled personnel. Zebrafish in the larval stage (72 hours postfertilization) can also be fin clipped[6][7] or sacrificed[8] for genotyping. However, due to their small size, fin clipping of larvae is technically challenging and time-consuming. Such health-invasive procedures can alter the swimming behavior of the fish, immune response, and more [9]. The discrete processing of individual fish also limits current genotyping, making it impractical for large-scale applications. Thus, there is a significant need for rapid, automatic, and non-destructive genotyping of zebrafish.

Some attempts have been made to utilize microfluidics to facilitate zebrafish genotyping. Samuel et al. developed a microfluidic system for fin clipping and collecting chorionic fluid for genotyping zebrafish embryos; however, it is unsuitable for high-throughput operations because the chip can only process one larva at a time. Moreover, the process requires continuous microscopic observation and manual pressure control for the movement of the larva inside the channel [10]. Venditti et al developed a Fin Scratching system to collect DNA from the tail of the zebrafish using a custom-made needle and apparatus to hold larvae [11]. The process involves manual steps of precise placement of larvae on a custom-made scratching support plate and scratching one larva at a time, making it a low-throughput system. Zhang et al used an enzymatic process for high-throughput genetic material collection from zebrafish [12] However, using pronase or proteinase K can affect the health of the embryos if they are not adequately washed immediately afterwards.

Recently, our lab developed an automated high-throughput device that can collect cells and usable DNA from zebrafish larvae for analysis by PCR, agarose gel electrophoresis, and HRMA (High-Resolution Melt Analysis) [13]. The Zebrafish Embryo Genotyper (ZEG) (Figure 1a) is a automated genotyping device consisting of a 3D-printed base unit and a chip that holds 24 zebrafish embryos in wells with a roughened bottom surface. A hydrophobic layer is used on the surface of the chip to create the wells and isolate aqueous media droplets. Finally, a cover is placed on top of the slide during device operation to limit evaporation. The chip suspended above the base unit is agitated at a specific frequency, creating an abrasive environment for the embryos that releases DNA.

**Figure 1.**
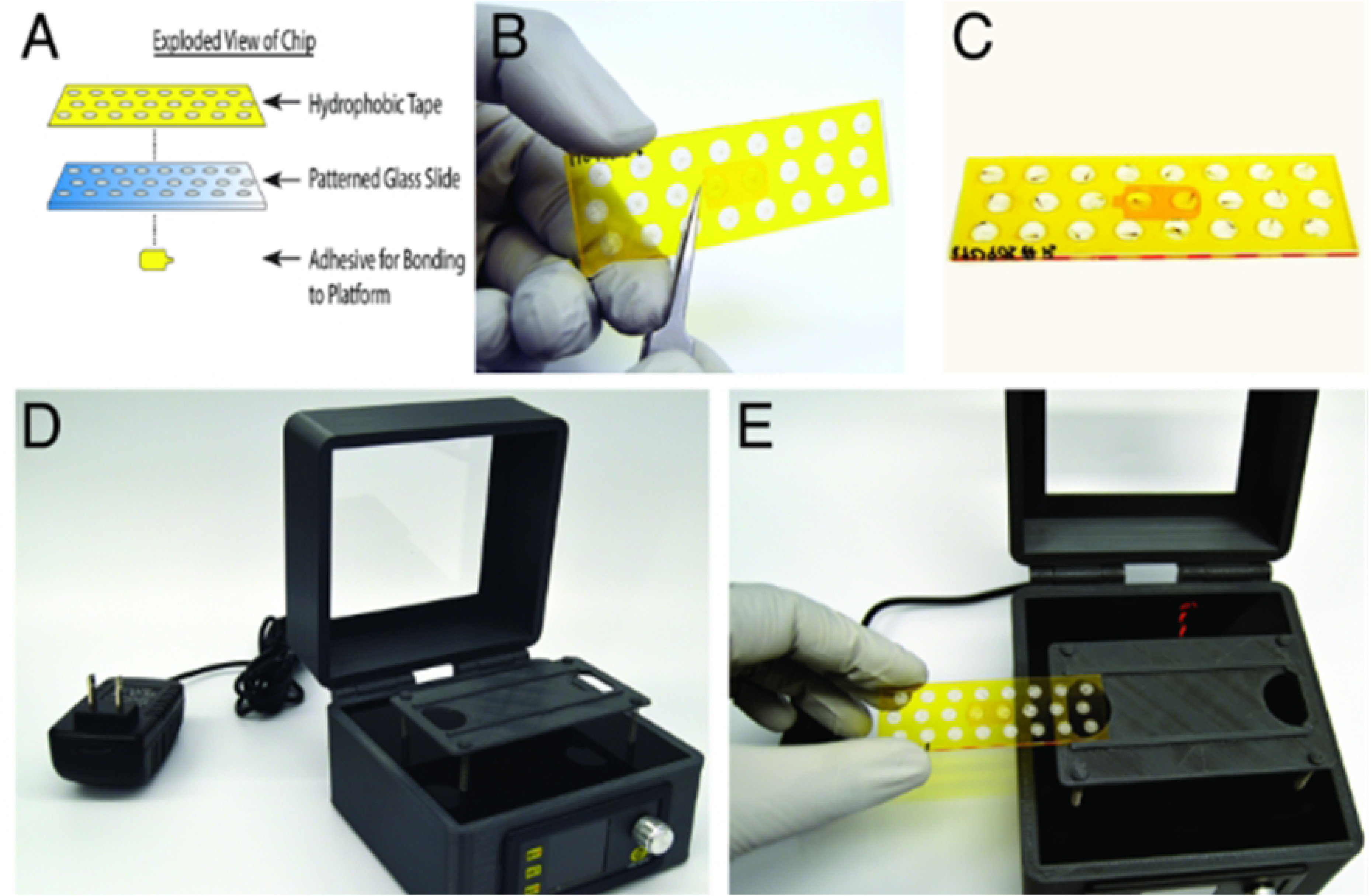
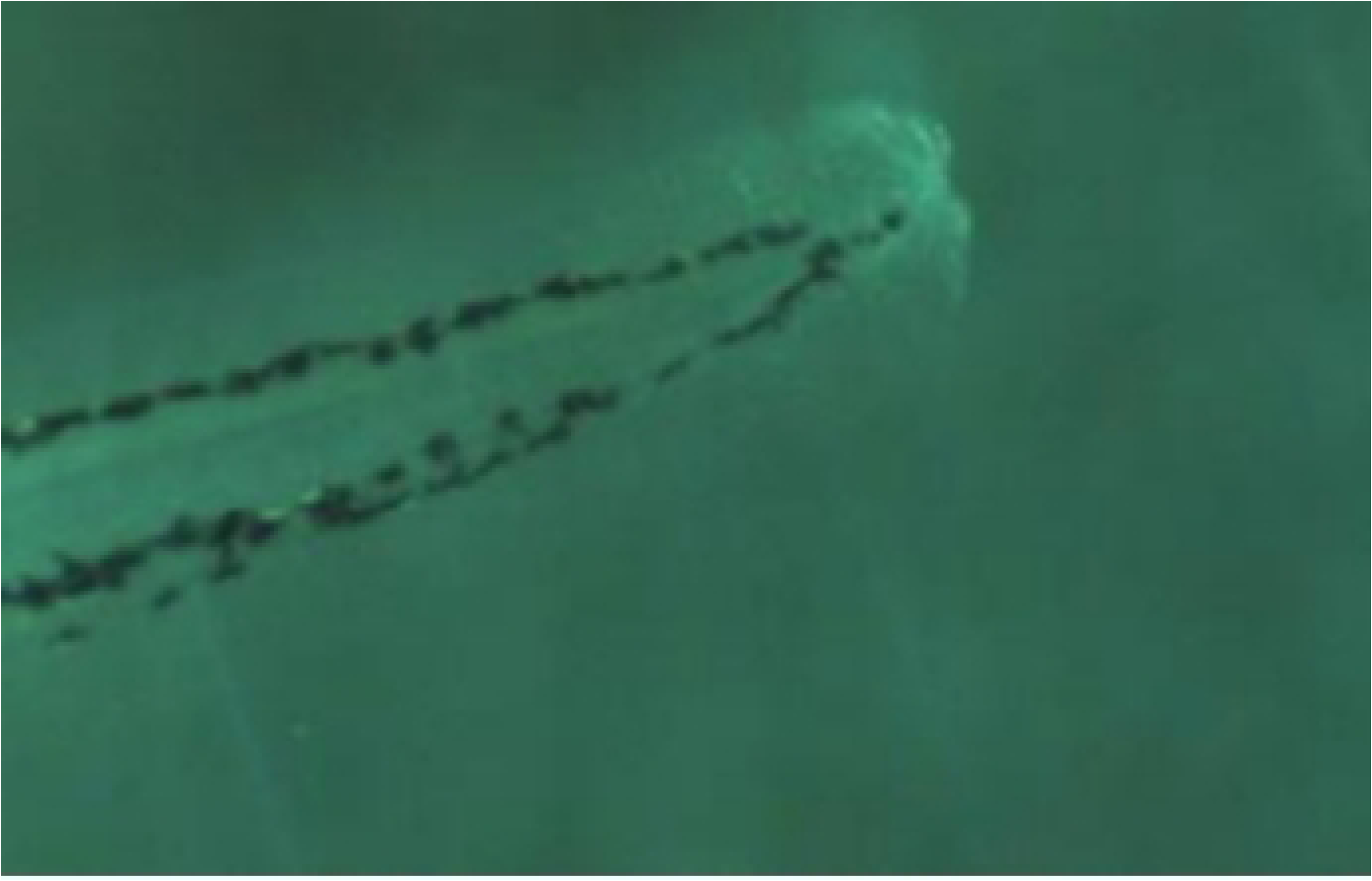
(a) ZEG device [13]- A) structure of the ZEG chip B) adhesive tape at the bottom C) ZEG chip with embryo loaded D) ZEG base unit E) ZEG chip loaded in base unit (b)the fin condition of the zebrafish embryo after ZEG operation

The ZEG device has been shown to effectively collect DNA from 48 to 72-hour post-fertilization (hpf) embryos, and the process can be performed simultaneously on 24 embryos. The extraction protocol takes 10 minutes to run, including loading and unloading of 24 embryos. The device requires minimal training and is simple to operate. Moreover, it is non-invasive to the embryos as it typically only scrapes small portions of the fin parts without harming the fish. The device has been shown to achieve a sensitivity of more than 90% while ensuring over 90% survival of the embryos [13]. The condition of the fin after the ZEG operation is shown in Figure 1(b). While these metrics are very good, the loss of even a few fish can be challenging in an experiment, so this work focuses on opportunities to improve the success of this approach and reduce fish and sensitivity losses.

Thus, in this work, we performed a design of experiments (DOE) analysis to evaluate the effects of overall chip surface chemistry, well surface roughness profile, and chip movement and loading parameters, including voltage, frequency, operation time, and loaded sample volume, on DNA collection using the ZEG. Quantitative polymerase chain reaction (qPCR) was used to quantify the DNA collection. Using the data generated, we performed a *t*-test and ANOVA analysis to determine the optimal parameters for operating the ZEG device.

## 2. Methods and Materials

### 2.1. Ethics Statement

The University of Utah Institutional Animal Care and Use Committee (IACUC) guidelines were maintained while performing the experiments on zebrafish, regulated under federal law (the Animal Welfare Act and Public Health Services Regulation Act) by the U.S. Department of Agriculture and Office of Laboratory Animal Welfare at NIH, and accredited by the Association for Assessment and Accreditation of Laboratory Care International. The University of Utah IACUC specifically approved this study

### 2.2. Fish Stock and Embryo Generation

Standard methods were followed for fish and embryo breeding. Embryos were raised in an E3 medium and maintained at a temperature of 28°C[14]. Fishlines used in this paper were Tg(myl7:EGFP; foxP2-enhancerA.2: Gal4-VP16_413-470_) ^zc72^ [15]; abcd1^sa509^, and abcd1^zc90^ [16]. Lines are available through the Zebrafish International Resource Center (ZIRC) or upon request.

### 2.3. ZEG Device Construction & Operation

The ZEG device consists of two significant parts: a 3D-printed (Prusa Mk3S+) base unit that contains a platform for placing disposable chips [13] and a locally roughened chip where embryos are loaded. The base unit can vibrate the platform at specific frequencies, creating an abrasive environment inside the wells on the chip; when the loaded embryo comes into contact with the rough surface, DNA is collected. The in-plane oscillation is created by suspending the platform on springs and attaching a vibrating motor (Precision Microdrive 312–108). The vibration motor’s frequency can be controlled by applying different voltages from a power supply (Uctronics U5168). An evaporation-limiting cover is also placed on top of the slide during the device’s operation.

The roughened chips used on the ZEG are disposable and, in this work, were made from standard microscope glass slides (75mm × 25mm). Each chip contains a total of 24 wells, arranged in three separate rows of 8. Laser etching was performed on the glass to create a rough, abrasive surface at the bottom of the circular wells. A double-sided hydrophobic polyimide tape with the same circular well configuration was cut out using a laser and applied on top of the slides and used to form the walls of the wells. The openings in the tape leave the roughened circles on the glass uncovered. Since the polyimide tape forming the walls of the wells is hydrophobic, when water droplets are placed in the wells, they form a dome shape. For some experiments, we applied a 3D-printed PETG layer that changes the shape of the liquid inside the well from convex to concave due to its hydrophilic nature, allowing more volume to be loaded.

In general, the device can be used for zebrafish embryos from 48 hpf to 7 days post-fertilization (dpf). In this paper, only 72 hpf embryos were considered for testing to eliminate any false negative results due to the varying embryo sizes at different ages. The 72-hpf embryos were loaded onto the chip wells with 8-12 µL E3 medium using a transfer pipette. The loading phase takes around 2-3 minutes. The chip was then placed on the device platform, and an evaporation cover was placed on top of it. The platform is normally agitated for 5 minutes, but the time can be varied from 3 to 10 minutes depending on needs. During this vibration phase, the embryos are forced to slide across the rough surface, resulting in scraping of the fin at the surface and the collection of cellular material. After the vibration phase, the E3 medium containing the scraped cellular debris was collected using a 10 µL pipette. The embryos were then transferred to 96-well plates with an E3 level of more than 300 µL. More details on the operation, component details, and flow diagram can be found in [13].

### 2.4. Optimization of the ZEG Device

#### 2.4.1. Optimization of device geometry

In this work, improvement of the ZEG chip was explored by modifying two characteristics: the roughness profile of the chip surface and the chip design. The previously developed roughness profile [13] was compared with a series of modified profiles to determine which profile provided the best DNA collection from the zebrafish embryos. The chip design was also modified to address the evaporation issue associated with the loaded water volume, and it is compared with the previous original design in DNA collection.

##### 2.4.1.1. Optimization of the roughness profile of the chip

The roughness profile of a ZEG chip is expected to play a significant role in determining the amount of DNA collected, so experiments were designed to test various profiles. To generate different profiles, the top surface of the glass slide was etched using a 75W CO_2_ laser system (Universal Laser Systems ULS 3.60) via a laser vector engraving technique, which was completed in approximately 11 minutes for the entire chip. The high heat condition from the laser causes temperature buildup in the chip, creating cracks and sharp edges. This creates an uneven surface texture or surface roughness. The power, speed, and image density settings of the laser system can be varied to create different roughness profiles. The power settings of the laser system range from 0% to 100% and modify to the depth of cut or engraving on the target material per pass; given a constant speed, a higher power setting provides a deeper cut. The speed setting can be adjusted from 0% to 100%, which accelerates or decelerates the system and consequently affects the cutting depth.

Various roughness profiles can be created on the glass surface by adjusting the power and speed in different combinations. Higher power and slower speed produce a deeper cut, while lower power and higher speed create a shallower cut. To simplify analysis, the power was held constant while varying the speed during engraving to create different roughness profiles. The spacing of the laser rastering passes is determined by the “image density” setting as shown in Supplementary Table 1. Initial experiments with varying image density and focusing/defocuing of the laser beam showed an impact on DNA collection from the fin of zebrafish larvae. After some preliminary testing, two chip manufacturing processes were chosen for comparison: the “modified” chips with an image density of 3 and a focused laser at 55% power and the “original” chips with a image density of 5 and a defocused laser at 65% power. These two chips were selected due to different roughness profiles (Supplementary Figure 1), which was expected to lead to different levels of DNA collection.

##### 2.4.1.2. Performance measurement by fin collection rating

A series of experiments were performed to compare the effect of the two roughness profiles on embryo survival and health after ZEG operation. Embryos were processed on the ZEG as described in later sections and the embryos were immediately transferred to a 96-well plate for storage while the fin condition was observed under a dissecting microscope with the examiner blinded to the roughness profile of the chip. Based on the tail condition of the embryos, a rating scale from 1-10 was developed; a higher rating indicates higher fin abrasion, and a lower rating indicates less abrasion. A rating of 10 was given by the examiner when the embryo was dead, or the fin was completely removed. Ratings of 1-9 were given to fish that were alive after the ZEG operation. The fish was considered alive when the heartbeat could be observed under a microscope. The rating conditions are listed in Supplementary Table 2, and the associated images are presented in Supplementary Figure 2. The same examiner performed all the fin-clipping observations to ensure consistency in rating. This value was designed to quantify the amount of abrasion resulting from the use of the ZEG system.

##### 2.4.1.3. Surface chemistry of the chip design

In the existing chip (original chip), with wells formed using hydrophobic tape, the water or E3 droplets in the wells form a dome shape with a larva inside. If we increase the volume of media from 10 µL to 12 µL, we could collect more media containing DNA. However, upon increasing the volume, the water line at the middle of the wells moves away from the rough surface due to the dome-shaped droplet. This provides sufficient space to move the larva away from the rough surface, resulting in reduced abrasion and lower DNA collection overall. Therefore, increasing the volume does not provide better performance with the current hydrophobic tape design. To determine if a different droplet profile would improve DNA collection while allowing larger sample volumes, a hydrophilic layer was tested instead of using hydrophobic tape. The hydrophilic layer or tape creates a concave water surface. As a result, increasing the volume in the wells has a relatively small impact on the location of the embryo relative to the surface, potentially maintaining larva-surface contact and, consequently, DNA collection while allowing larger volumes to be used.

As part of this optimization effort, we 3D-printed a 1.5 mm thick middle layer containing holes using PETG. Figure 2 shows the schematic of the 3D-printed layer. For this modified design, only the original roughness profile of the glass slide was tested. An optical profiler (Olympus OLS5000-SAF) was used to analyze and quantify the water surface profiles. The original chip wells were loaded with 12 µL of water, and the modified chip wells were loaded with 15 µL. qPCR was performed on the collected DNA to evaluate the effects of various parameters and optimize the design.

**Figure 2.**
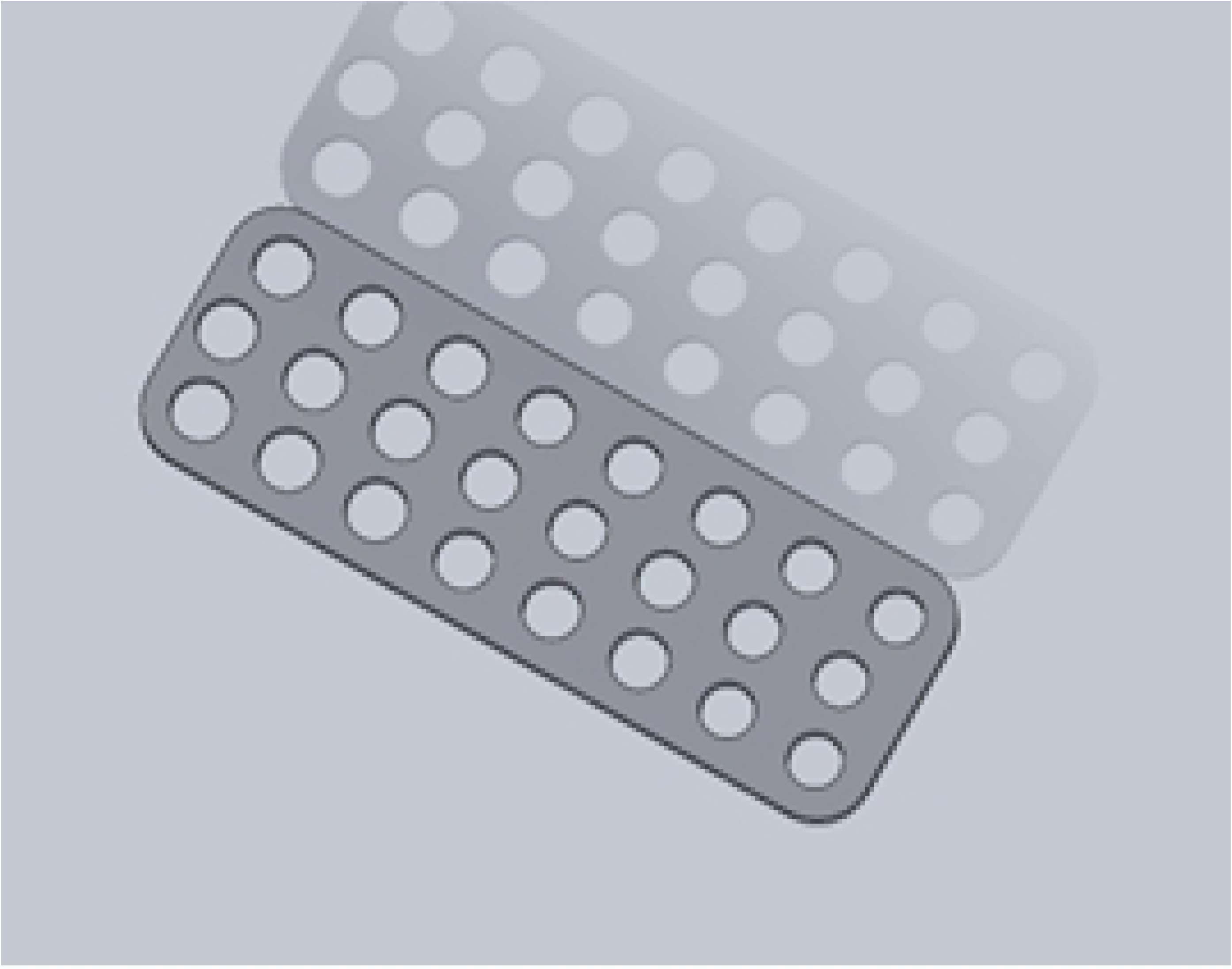
The design of the 3D printed layer.

#### 2.4.2. Parametric Optimization

Voltage and volume parameters are known to impact the performance of the ZEG device. Therefore, experiments varying voltage (1.4 V, 1.9 V, and 2.4 V) and volume (8 µL, 10 µL, and 12 µL) were performed. The voltage was applied continuously for 5 minutes. Each experiment was repeated *n* = 24 times with three different chips to measure the consistency of the results.

1.4 V, 12 µL volume, and 5 min continuous motor running were considered for the original design. In the modified design, we considered 1.9V and 2.4V to determine the effect of vibration frequency increases on DNA collection. To alleviate evaporation during loading, an increase in volume was necessary during this optimization. So we considered 15 µl and 20 µl for the proposed design.

Moreover, time and on/off operation could be another parameter that affects DNA collection, which has not been previously tested. Therefore, instead of running the ZEG device continuously for the 5 minutes of operation, we tested on/off operation. Here, we have considered the following durations: 5 minutes continuous, 5 minutes with 30-second on/off intervals, 5 minutes with 15-second on/off intervals, and 5 minutes with 5-second on/off intervals.

### 2.5. Quantification of DNA Collection

qPCR was performed to measure the DNA collected from each well of the ZEG chip. The targeted gene was abcd1 of zebrafish, which is highly conserved. GoTaq Hot Start Colorless Master Mix, Promega was used for genotyping, with the following primers and conditions: (forward primer) 5’-AGGTTGGCAAACCCTGACCA-3’ and (reverse primer) 5’-GTGTTGGCGCCTTTGGATTC-3’; 95°C for 3 min (m), followed by 40 cycles of 95°C for 15 sec (s), 60°C for 15s, 72°C for the 30s, and a final extension of 72°C for 2m.

The DNA collected through the manual fin clipping process was considered the positive control, and the DNA amount was measured using a Nanodrop. The positive control contained a DNA concentration of 172 ng/µL. Five dilutions (1/10, 1/100, 1/1000, 1/10k, 1/100k) were created from the positive control, and qPCR was performed for each run. The qPCR provided Ct values, which were used to create a standard curve with known positive dilutions. The standard curve and Ct values of the samples collected from the wells during the qPCR provided an indication of the amount of DNA collected. For the negative control, E3 media and TE media were used without any added DNA. Moreover, E3 media was placed inside the well and then collected as a negative control to verify whether any noise in the wells resulted in false-negative qPCR results. The qPCR was performed while optimizing the parameters of the modified chip design and simultaneously comparing them with those of the original chip design.

### 2.6. Measurement of Vibration Frequency

In the ZEG device, the voltage applied to the platform is directly associated with the motor’s vibration frequency. So, increasing the voltage should increase the frequency, hence creating more abrasion to the fin of the larva. A 3D laser vibrometer and oscilloscope were used for the measurements of the vibration frequency in correlation with applied voltage.

### 2.7. Survival and Morphological Analyses

The examiner who conducted the fin clipping observation performed survival and morphological analysis under a dissecting microscope. The observation was completed just after using the device and 24 hours after the experiment. The larvae were considered to have survived when their heartbeat was regular. The experiment was repeated multiple times with more than 25 embryos each time.

### 2.8. Statistical Analysis

Statistical analysis was performed using JMP (SAS), and a two-way t-test was conducted for comparison. ANOVA with post hoc Tukey’s HSD was performed to compare individual means across multiple groups and interaction profiles.

## 3. Results & Discussion

### 3.1. Optimization of the ZEG Device

#### 3.1.1. Characterization of the Roughness Profile

The surface roughness profiles of modified chip and original chip are presented in Figure 3(a) and 3(b), respectively, and the profilometer output is presented in Supplementary Figure 1(a) and 1(b), respectively. The surface profile shows that for focused laser beam with modified chip profile, the waviness is uniform and the number of waves for is much higher (pitch greatly reduced) compared to original chip roughness profile. However, the original chip, which used a defocused laser beam, does not exhibit uniform roughness. The focused laser beam targets a reduced area, and the defocused beam targets a larger area, though the depth of cut is about 5 µm deeper. The surface roughness, visible under a dissecting microscope, is shown in Supplementary Figure 1(c).

**Figure 3.**
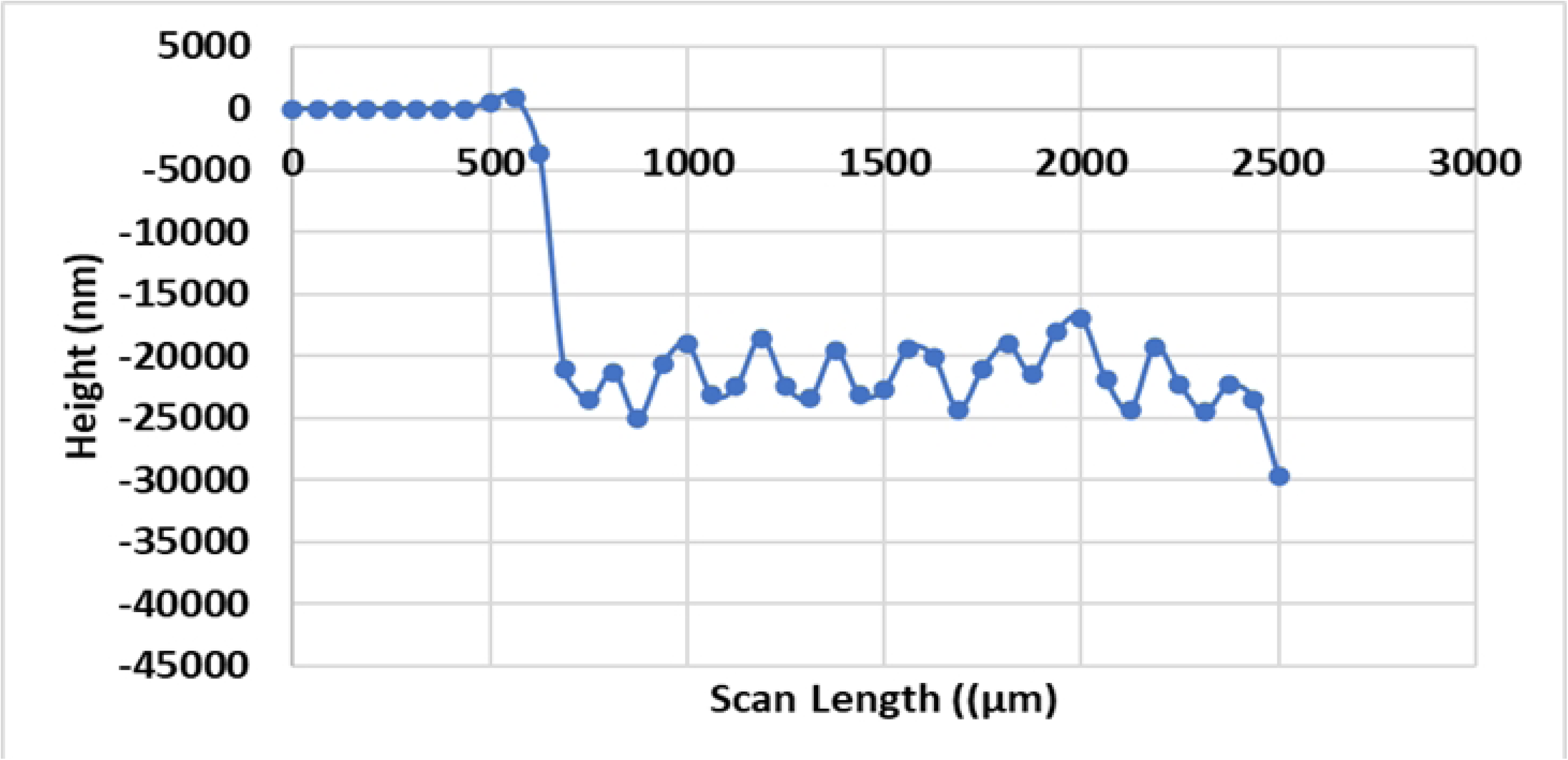
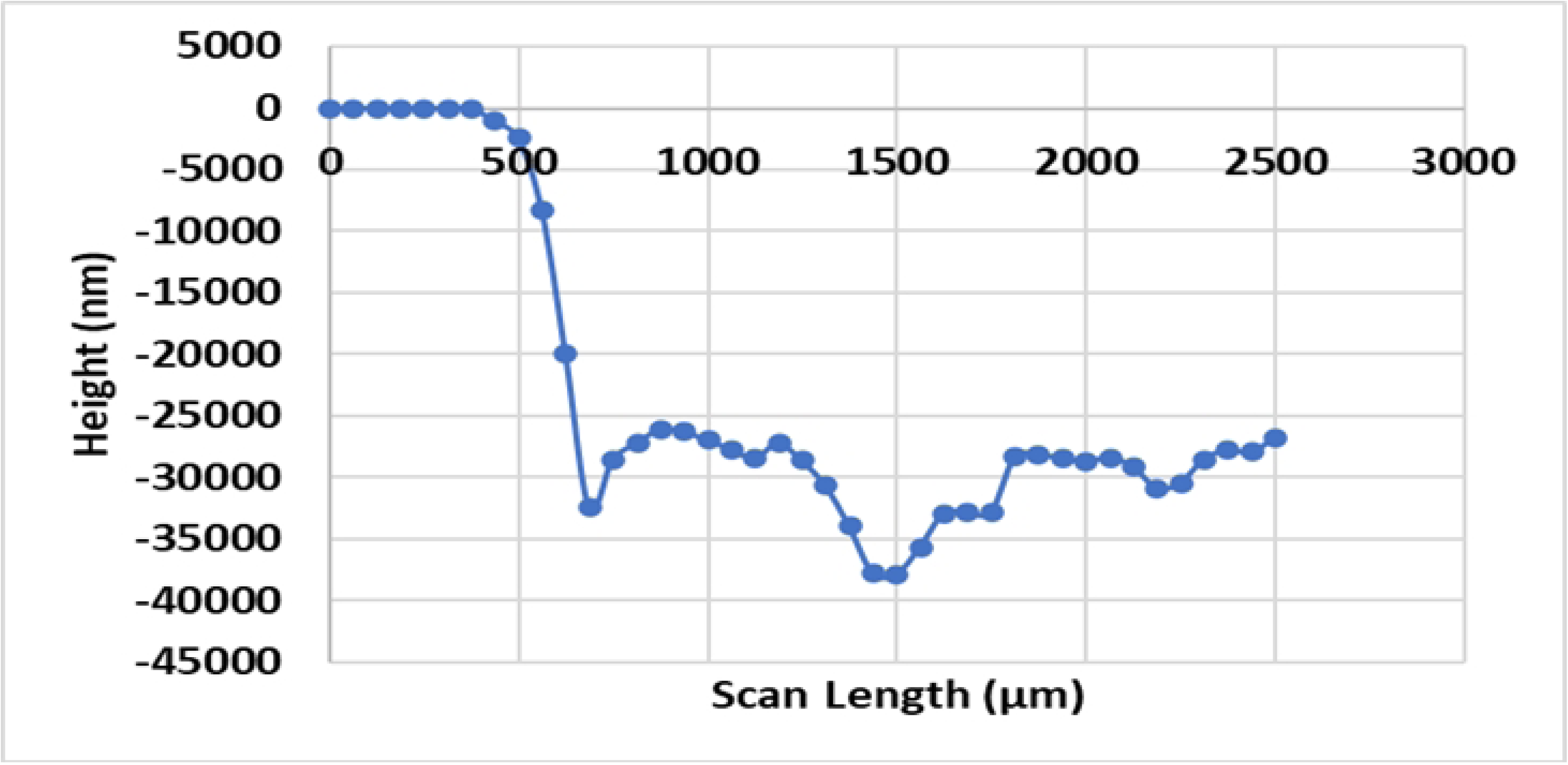
The roughness profile of the ZEG chip (a) Surface profile of modified chip (b) Surface profile of original chip

#### 3.1.2. Optimization of the Chip Design

The water surface profile difference resulting from the original chip and the proposed modified chip with 3D printed middle layer can be visualized in Figure 4(a). Figures 4(b) and 4(c) compare the water surface profile outputs resulting from the original and the modified design, measured using the optical profiler. The optical profiler output is shown in Supplementary Figure 3. The contact angles for the original and modified chips were measured to be 6.31° and -8.25° (ImageJ), respectively. The proposed modified design was able to accommodate 15 µL of water compared to 12 µL in the original design with a hydrophobic middle layer. This validates the idea behind the chip design change (from a hydrophobic middle layer to a hydrophilic middle layer), which was intended for collecting a larger volume of media from the chip well. Collection of a larger volume simplifies further PCR analysis without compromising the surface contact issue with the larva during the ZEG operation.

**Figure 4.**
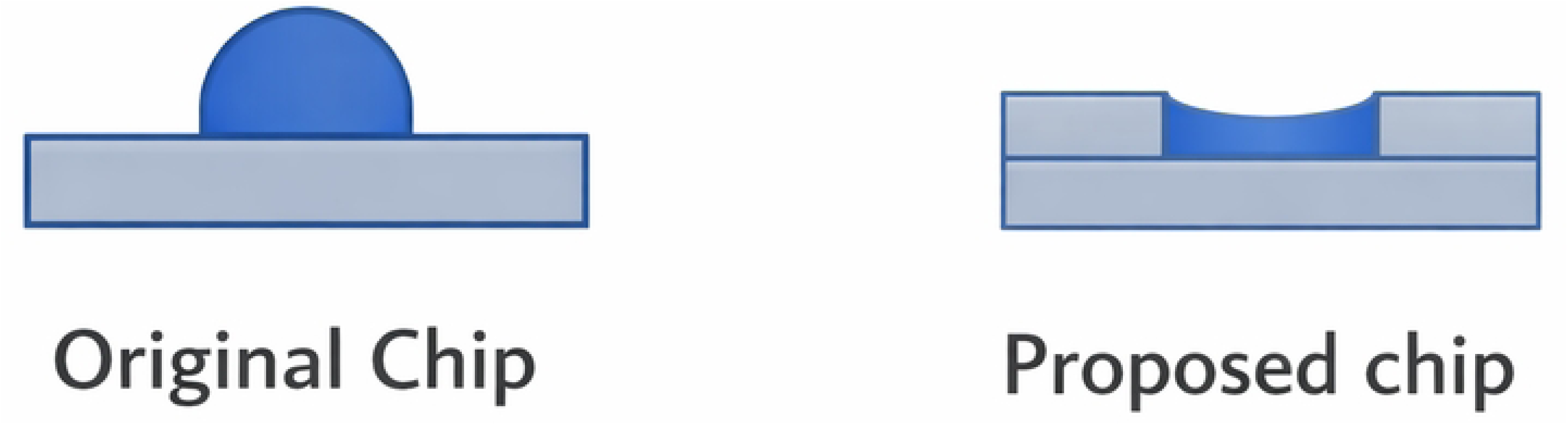
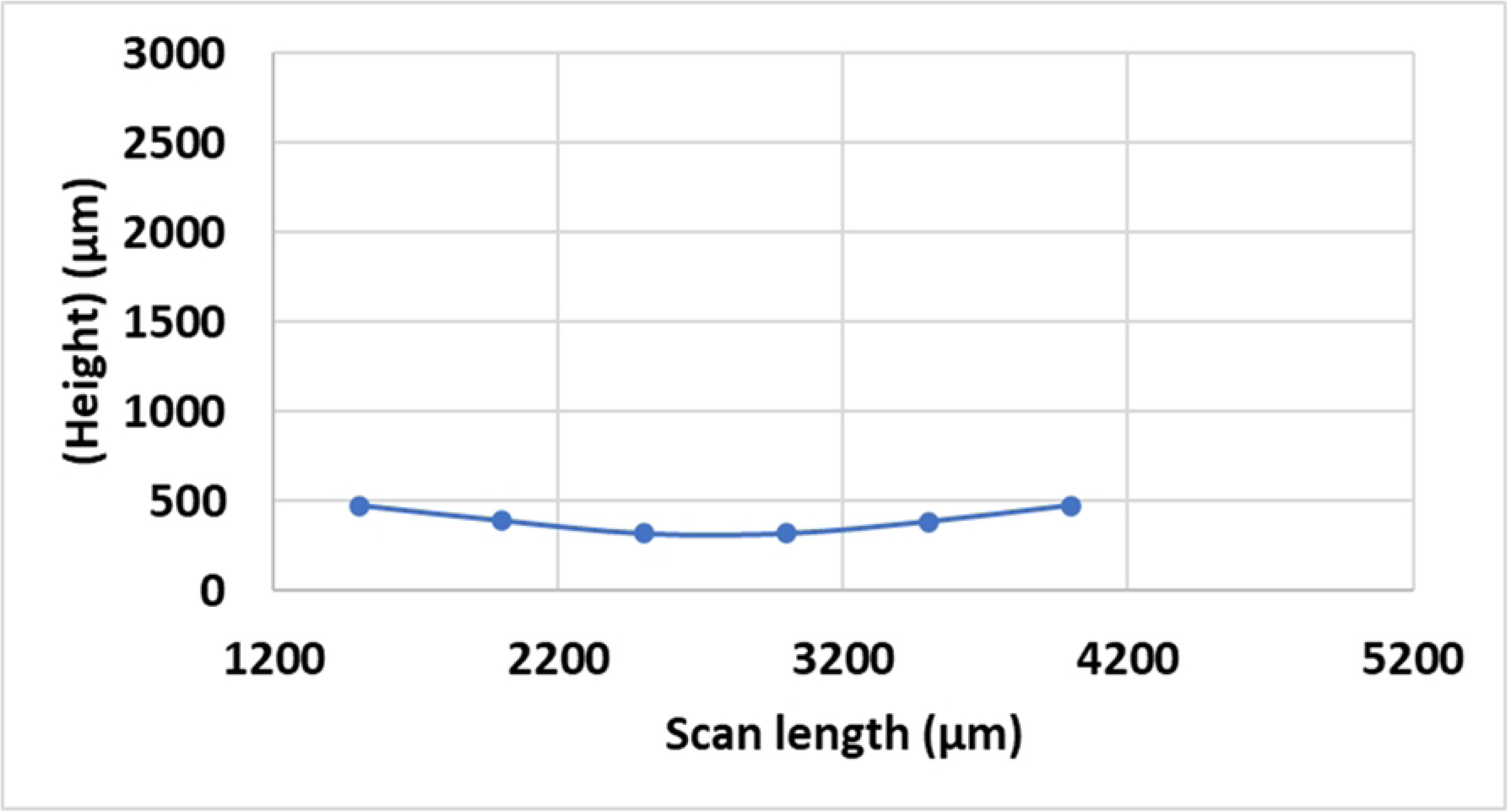
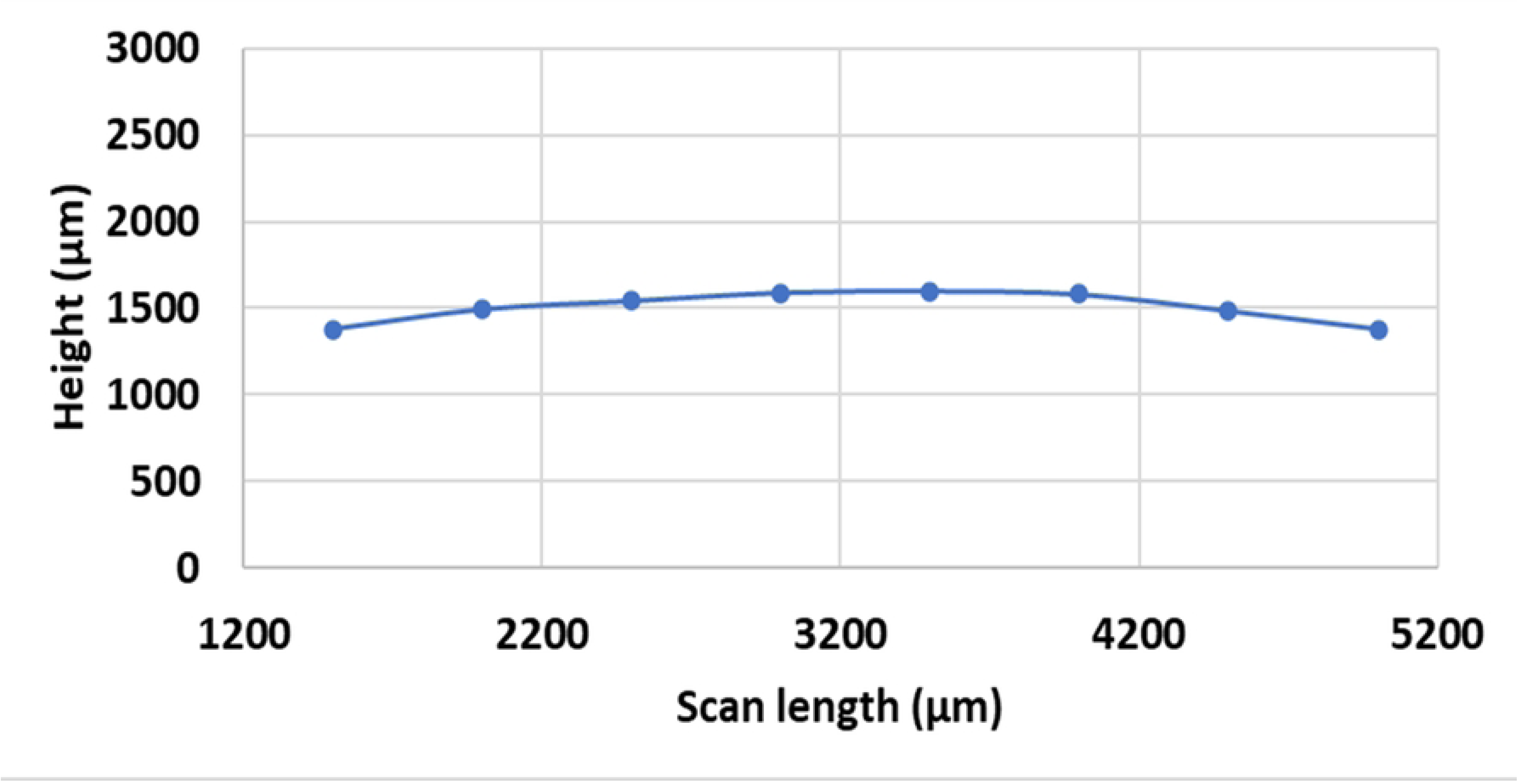
(a) Difference in well profile for hydrophobic compared to hydrophillic surfaces. The original hydrophobic chip creates a convex droplet, while the proposed modified hydrophilic design generates a concave shape. (b) Water surface profile for the original chip with 12 µL of a water droplet in the well. (c) Water surface profile of the proposed modified chip after loading a 15 µL water droplet in the well.

#### 3.1.3. Characterization of vibration frequency

The lateral vibration frequencies in the platform’s *X* and *Y* directions, resulting from applying different voltages to the system, are presented in Supplementary Table 3. For 2.4 V, the vibration frequencies in the *X* and *Y* directions are 212.8 Hz and 202 Hz, respectively. However, for 1.4 V, the frequency is lower (119.1 Hz in the X-direction and 117.6 Hz in the Y-direction). From the measurements, it can be validated that the vibration frequency is higher for high-voltage applications and lower for low-voltage applications.

### 3.2. Optimization Based on Fin-Condition Ratings

#### 3.2.1. Roughness Profile Effect on Fin-Condition Ratings

The fin condition ratings after ZEG processing are expected to correlate with the amount of DNA collected during the process. Specifically, higher fin-condition ratings should indicate that more material was removed from the fish and more DNA collected. The measurements of the cellular material or DNA collection as a function of roughness profile is presented in Figure 5. The collected data indicates fin collection in for different chips and operating conditions. specifically, Figure 5(a) shows the measured of fin condition observed for 1.4 V at three different volumes for R3F and R5DF chips. It shows that 8 µL provides the highest level of fin collection (fin rating = 5), compared to 10 µL (fin rating = 4.5) and 12 µL (fin rating = 4.3).

**Figure 5.**
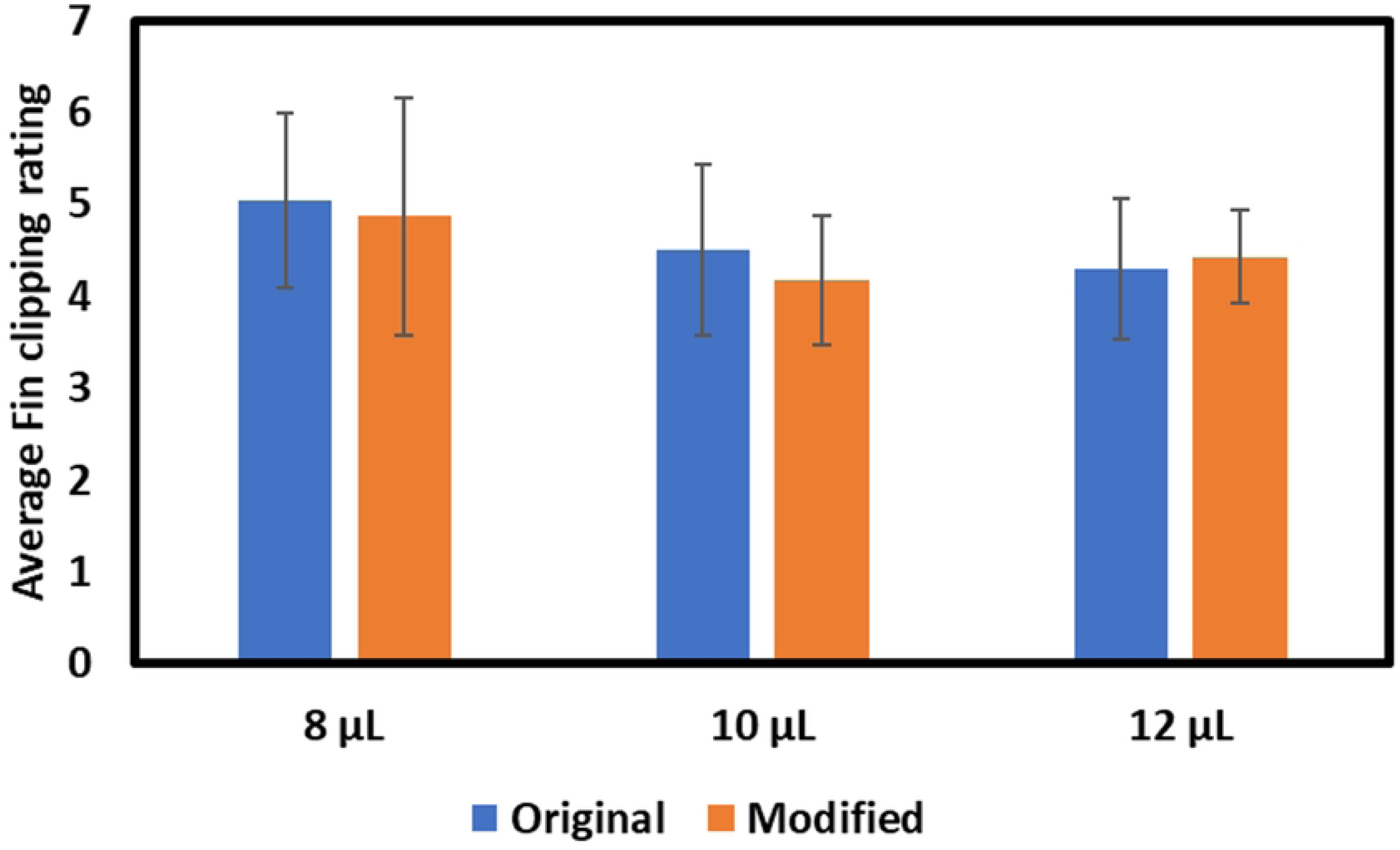
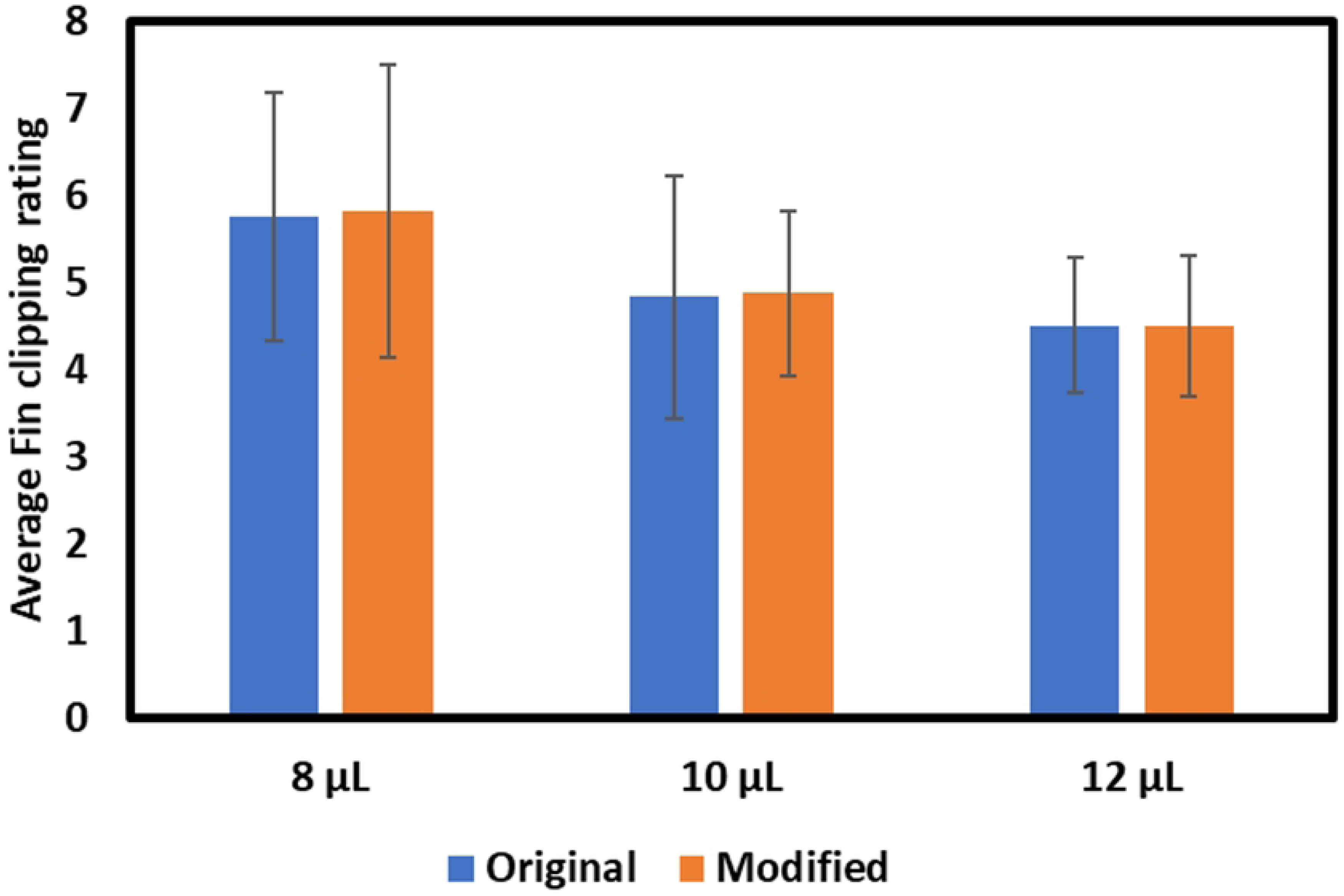
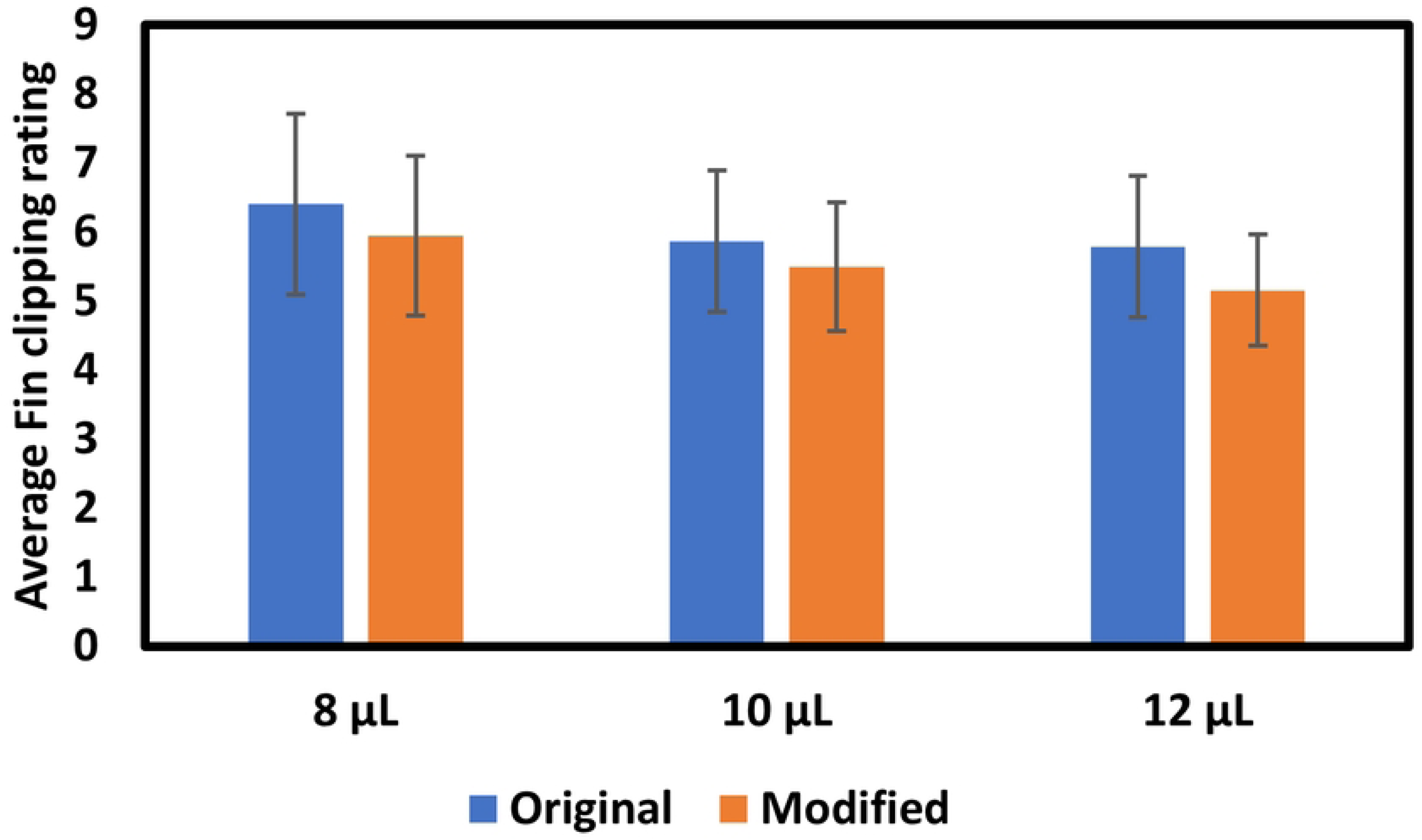
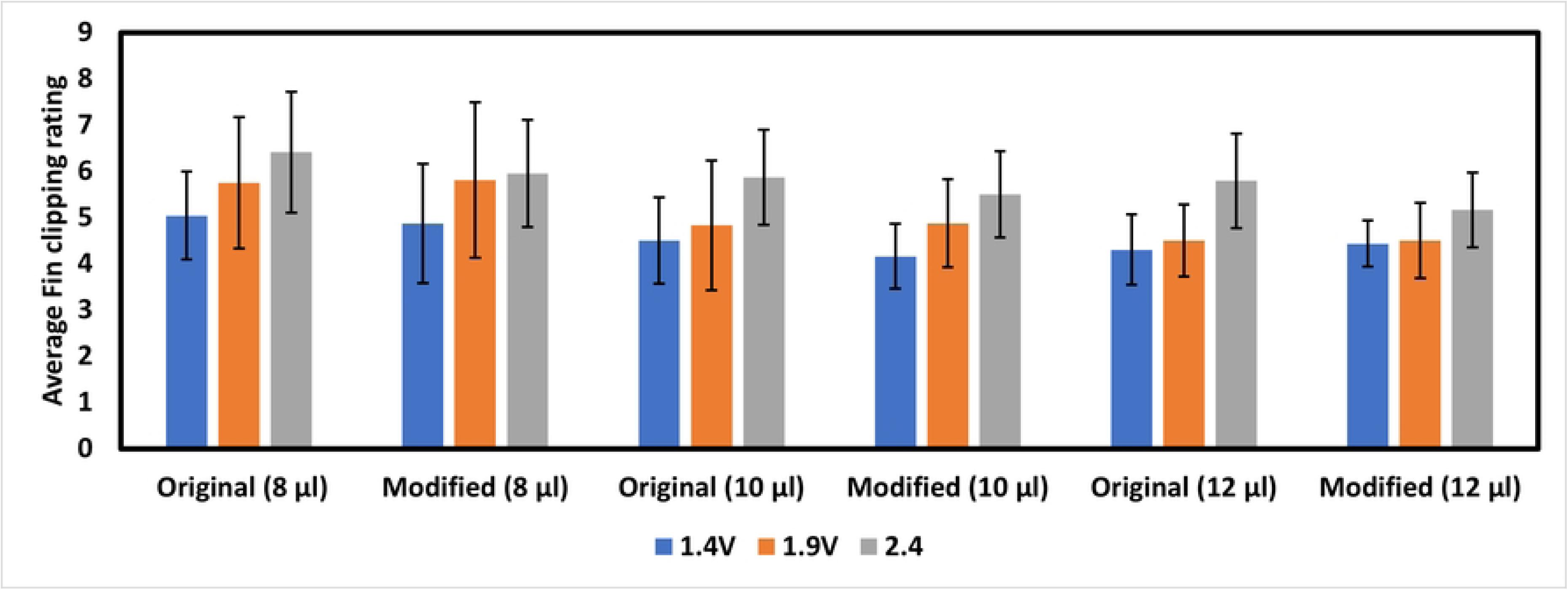
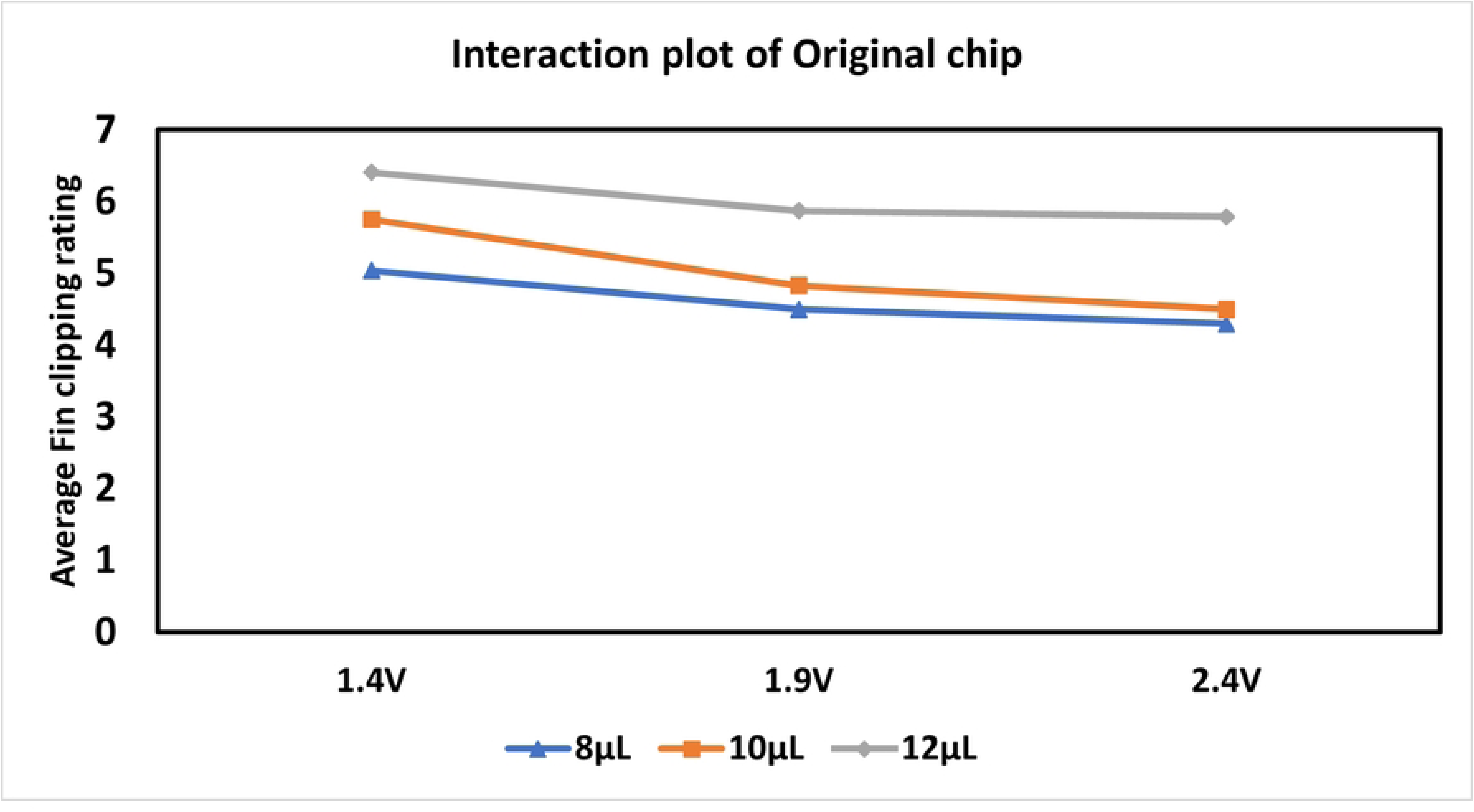
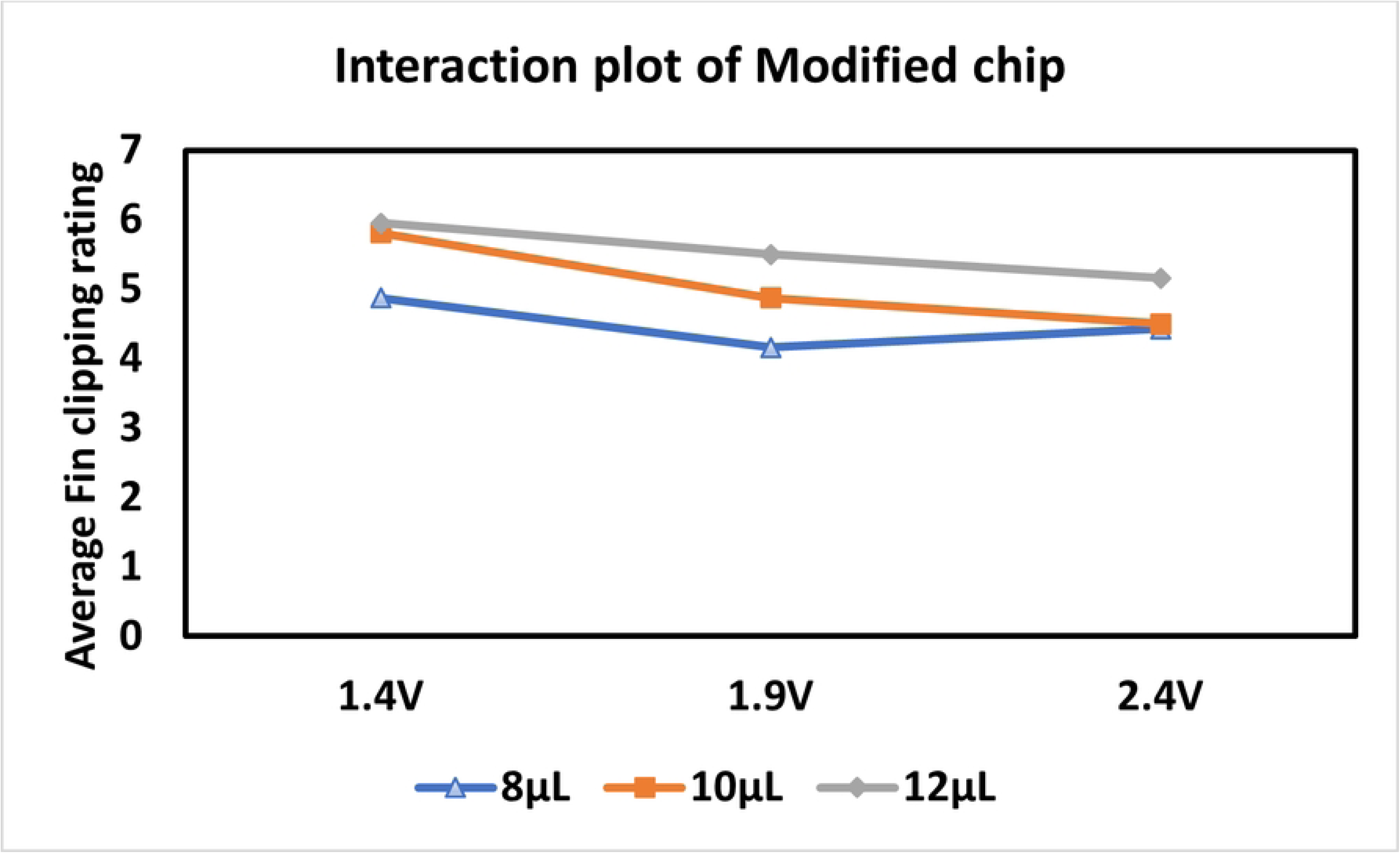
Fin-clipping rating for different conditions. Comparison of original and modified roughness for the average fin-collection rating based on different volumes when (a) 1.4 V applied to the system, (b) 1.9 V applied to the system, and (c) 2.4 V applied to the system. (d) Overall comparison for different voltages on roughness profile of original and modified chip. (e) Interaction profile for original profile (f) Interaction profile for modified profile

Moreover, comparing the average ratings for the roughness profiles shows that original collects more cellular material than modified profile for different volumes. A similar trend can be observed in Figures 5(b) and 5(c), where the fin collection levels are presented for 1.9 V and 2.4 V, respectively. Here, 8 µL again provides the highest level of fin collection, with values of 5.75 and 6.41 for 1.9 V and 2.4 V, respectively. However, at 1.9 V (Figure 5(b)), the fin condition ratings for the different roughness profiles at each volume were visually almost negligible and statistically not significant (p>0.05).

Figure 5(d) shows the overall rating of fin collection for different volumes and voltages for the roughness profile of original and modified chip. From the figure, it can be observed that the higher frequency provides a higher level of fin collection. Also, 2.4 V provides a higher fin collection for all roughness profiles and volumes.

Interaction plots are generated to visualize the combined effect of volume, voltage, and roughness profile and are presented in Figure 5(e) for original profile and Figure 5(f) for modified profile, respectively. For both modified and original roughness profiles, the trend is largely parallel across volumes, indicating weak interaction between voltage and volume. For modified profile, fin rating decreases from 1.4V to 1.9V across volume followed by minimum changes for 2.4 indicating limited interaction with non-monotonic voltage response. Whereas original shows a monotonic decrease in rating with an increase in voltage across volume. Overall, voltage and volume act as additive main effects, with roughness mainly affecting how consistently fin clipping rating changes with voltage. The quantification and statistical significance of these main and interaction effects are discussed in the following section.

#### 3.2.2. Statistical Analysis of Fin-Collection Ratings

A two-way ANOVA was performed to determine the main and interaction effects of roughness profile, voltage, and volume on fin-collection rating. The analysis revealed no statistically significant interaction between the effects of roughness profile and volume (F = 0.1916, p = 0.66). Similarly, no statistically significant interaction was found when voltage and roughness profile were considered (F = 1.82, p = 0.177), nor was there a statistically significant interaction between voltage and volume (F = 0.163, p = 0.68).

Simple main effect analysis revealed that the roughness profile statistically affects the fin collection rating (p = 0.0355). Moreover, the main effect analysis also revealed statistical significance for volume (p < 0.0001) and voltage (p < 0.0001). It indicates that different roughness levels, voltage, and volume affect the fin collection after ZEG operation. The interaction profile and effect test data are presented in Supplementary Figures 3 and 4.

A two-sample *t*-test was performed to compare two different levels of roughness, original and modified. The null hypothesis was that the mean fin collection for two roughness profiles is the same, and any observed difference between the sample means is due to natural sampling variation (chance). In the analysis for the null hypothesis, p = 0.07 indicating no significant difference between the performance of modified and original when considering the fin collection rate.

However, for a one-tailed *t*-test, we found that the mean of roughness original profile can significantly improve the fin-collection rating (p = 0.0351) compared to modified profile, considering α = 0.05. The analysis of the *t*-test, considering unequal variance, is presented in Supplementary Figure 5.

Prior to performing the ANOVA, assumptions of normality and homogeneity of variance were evaluated. The Shapiro-Wilk test applied to the ANOVA residuals indicates a deviation from normality (p<0.05), which is expected for large datasets and does not invalidate the analysis. Using Bartlett’s and Levene’s tests, variance homogeneity was examined. Though Bartlett’s test suggested unequal variances (p<0.05), Levene’s test showed no significant difference in variances across groups (p>0.05). Given the robustness of ANOVA to mild departures from normality and variance homogeneity, the ANOVA results remain reliable. The test data are presented in Supplementary Figure 6.

#### 3.2.3. Survival, Development, and Behavior After ZEG Operation

During ZEG operation, the embryos are exposed to mechanical vibration, and tissues are collected due to their contact with the chip’s rough surface, which could potentially damage the fish. Therefore, it is crucial to determine whether the ZEG operation affects the embryos’ natural behavior, development, and survival. The embryos were kept in an incubator with a temperature of 28°C for 24 hours, and survival was monitored based on heartbeat and movement. The survival rate of the embryos after the fin collection operation for different volumes, chips, and voltages is shown in Figure 6. The figure shows 100% survival for lower voltages and more than 95% for voltage 2.4. Statistically, the *p*-value is higher than the significance level (α = 0.05) for all conditions, indicating that survival at higher voltages is not significantly different from that at lower voltages. Higher voltage shows some bloated states in less than 5% of fish, which does not create a statistically significant difference in survival.

**Figure 6.**
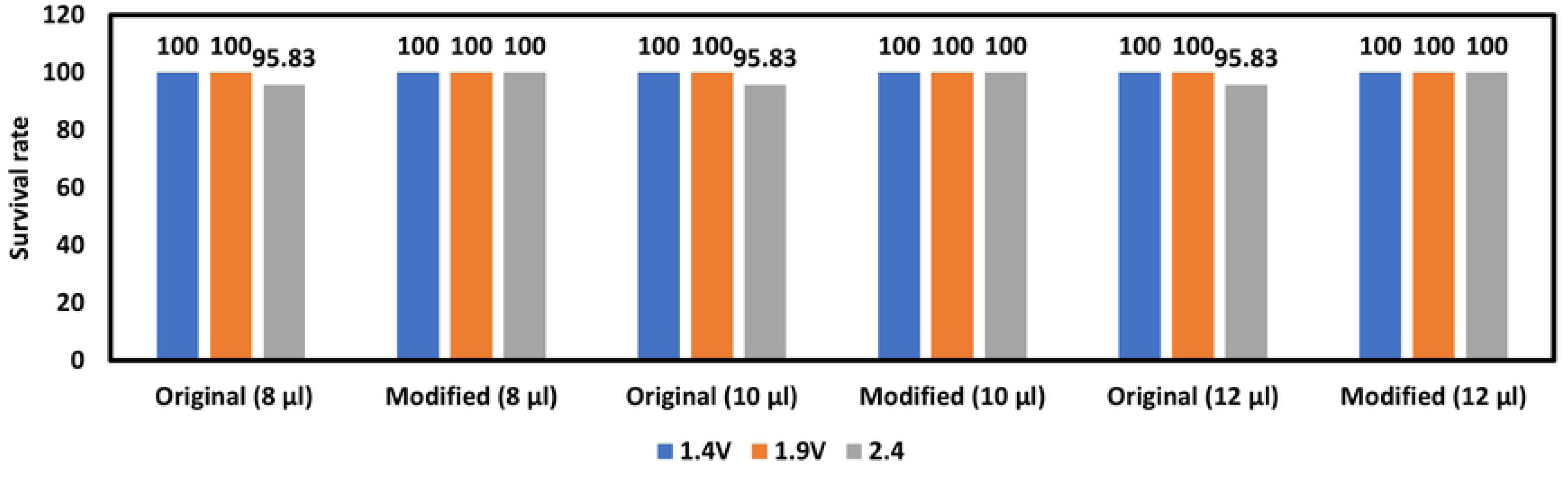
The survival rate of embryos after fin clipping for voltages: 1.4 V, 1.9 V, and 2.4 V; volume: 8 µL, 10 µL, and 12 µL; and roughness: R5DF and R3F.

### 3.3. Optimization of DNA Collection

#### 3.3.1. Effect of the Design Change

The concentration of DNA collected on the ZEG chips depends on the mass of the DNA released and the volume in the well. Generally, speaking, the goal was to maximize this concentration while providing sufficient volume for qPCR analysis . Figure 7 shows the average DNA collected (pg/µl) for different ZEG operating parameters. For each parameter, *n* = 6 samples were considered. Compared to the original design, we have identified three conditions based on the results shown in Figure 8 for the modified chip, where the average DNA collection is increased. Those conditions are: i) 2.4 V, 15 µL, 5 min 30 seconds on/off, ii) 1.9 V, 20 µL, 5 min 5 seconds on/off, and iii) 2.4 V, 15 µL, 5 min 30 seconds on/off. However, when total DNA collection was considered, we observed ten samples with values higher than the previous approach (15.15 pg). That is because the modified chips are loading higher volumes (15 µL and 20 µL) than the original chip. Therefore, if we consider average DNA collection, then it can be observed that the modified chip for 15 µL volume, 2.4 V, and 5 minutes running of the ZEG device with 5 seconds on/off provides the best average DNA collection (2.68 pg/µl) compared to other conditions and the original chip (1.64 pg/µl). Even when optimizing for total DNA, including the total volume from the well, it provides the highest total amount of DNA mass (38.9 pg) for each data point presented.

**Figure 7.**
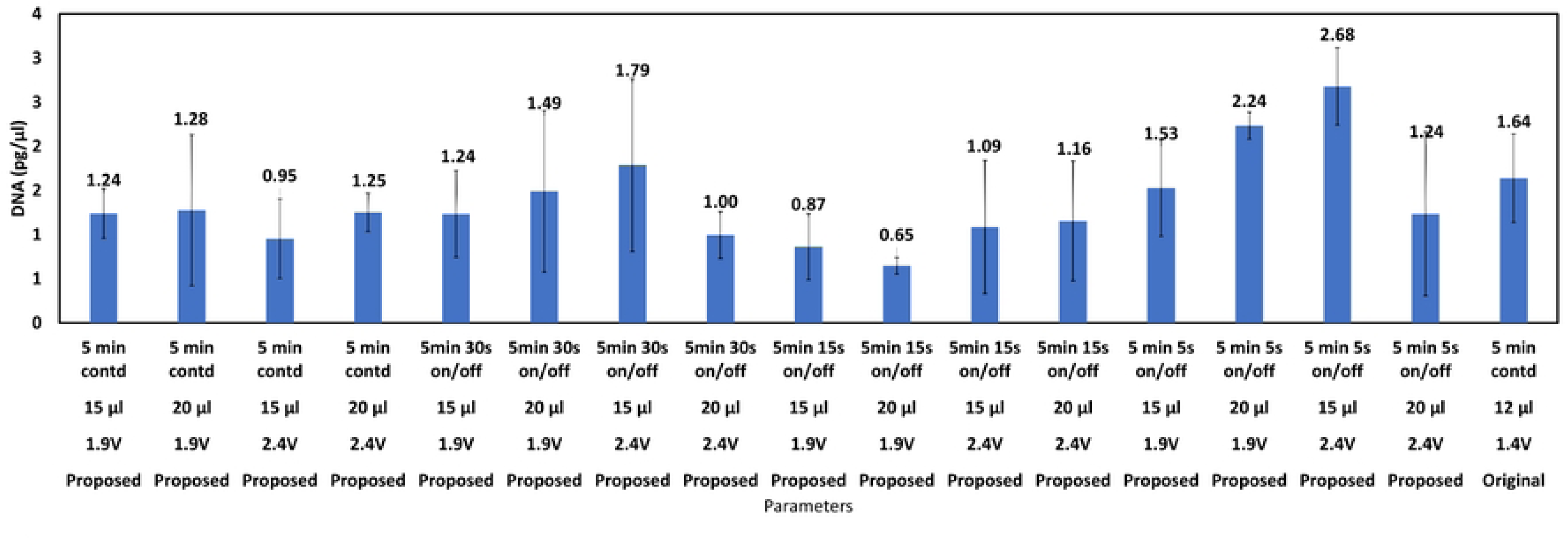
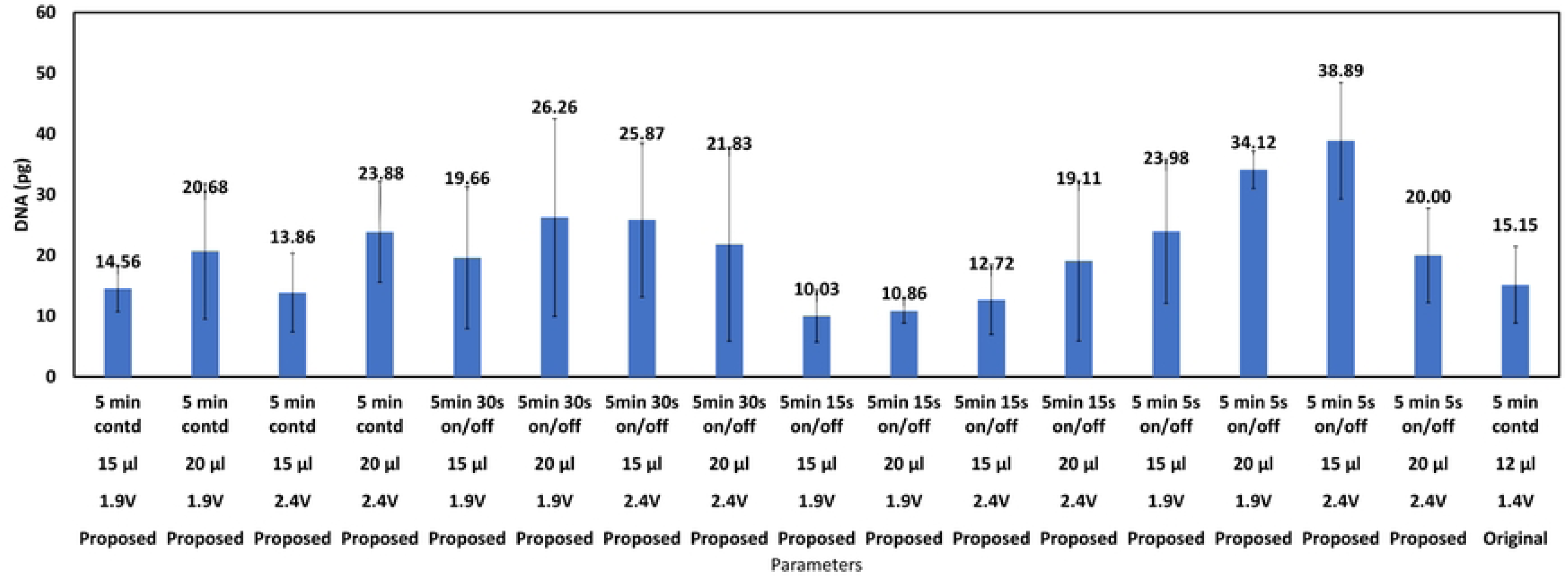
(a) Average DNA collection (pg/µl) for different values of parameters - voltage, droplet volume, and time of ZEG operation. (b) Total average DNA collection (pg) for different parameters. 6 samples were included for the average calculations.

**Figure 8.**
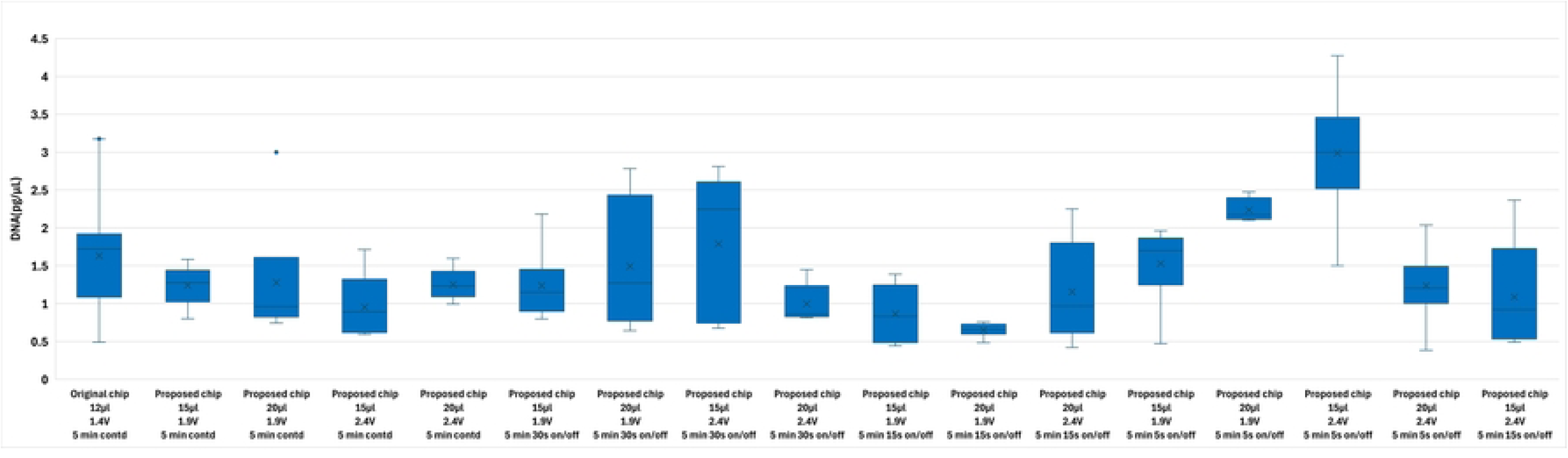
Box and Whisker plot of the average DNA collection (pg/µl) for different values of parameters - voltage, droplet volume, and time of ZEG running.

The average DNA collection (pg/µL) for different voltage, droplet volume, and time parameters is presented in Figure 8. The figure shows that the proposed modified chip design, combined with the parameters of 15 µL, 2.4 V, and a 5-minute running time with a 5-second on/off cycle, provides the best results.

Now, the proposed modified chip, with parameters of 15 µL, 2.4 V, and a 5-minute running time with 5 seconds on/off, was compared with lower volumes of 12 µL and 10 µL, while keeping other parameters constant. Figure 9(a) presents the comparison of average DNA collection (pg/uL), and Figure 9(b) shows the total DNA collection (pg) considering the volume. An original chip with 12 µL, 1.4 V, and 5-minute continuous running is also compared. The figure shows that a lower volume with a proposed modified chip provides better performance than the original chip for both conditions. The modified chip with a 15 μL volume performed better when the total DNA amount was taken into account. However, when the average DNA concentration (pg/µL) was considered, the 10 µL also performed better than the 15 µL.

**Figure 9.**
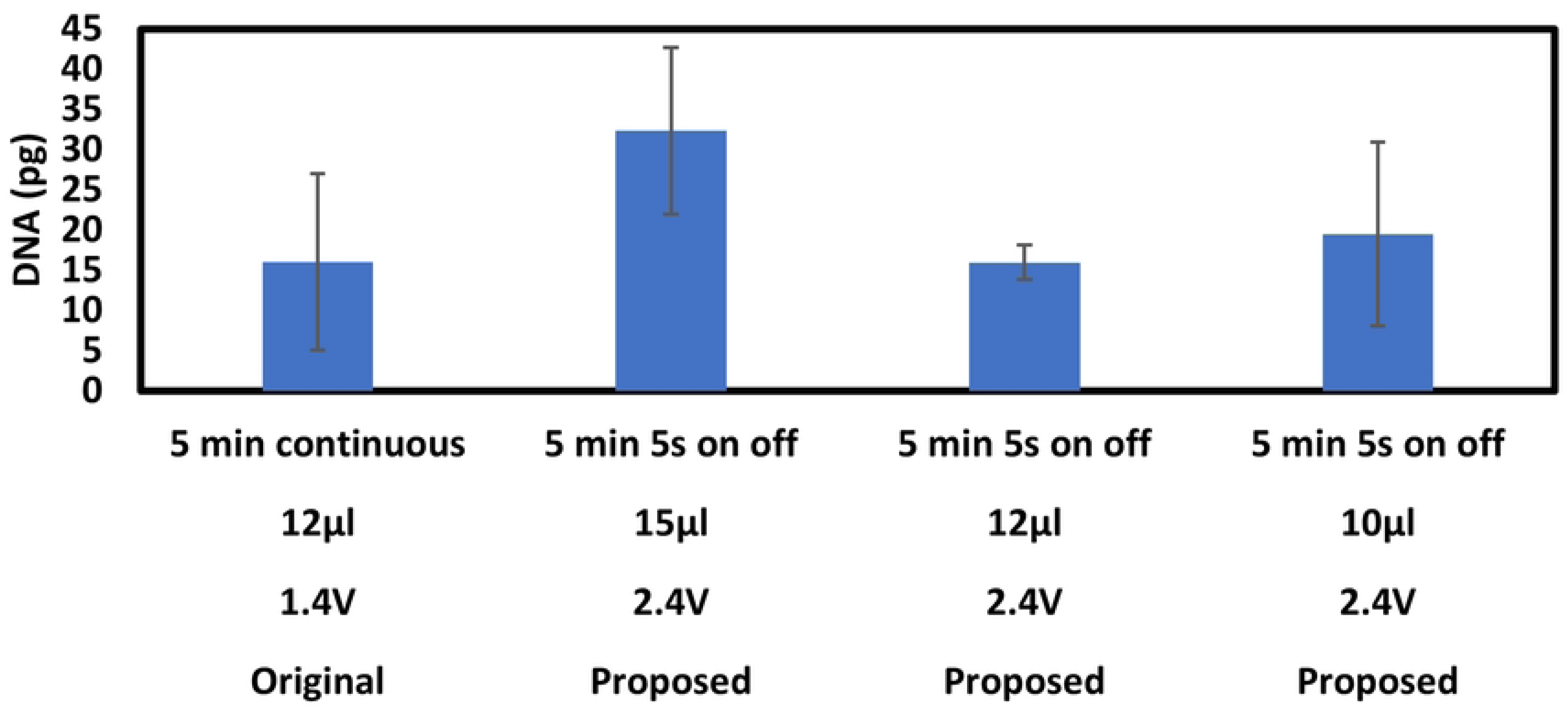
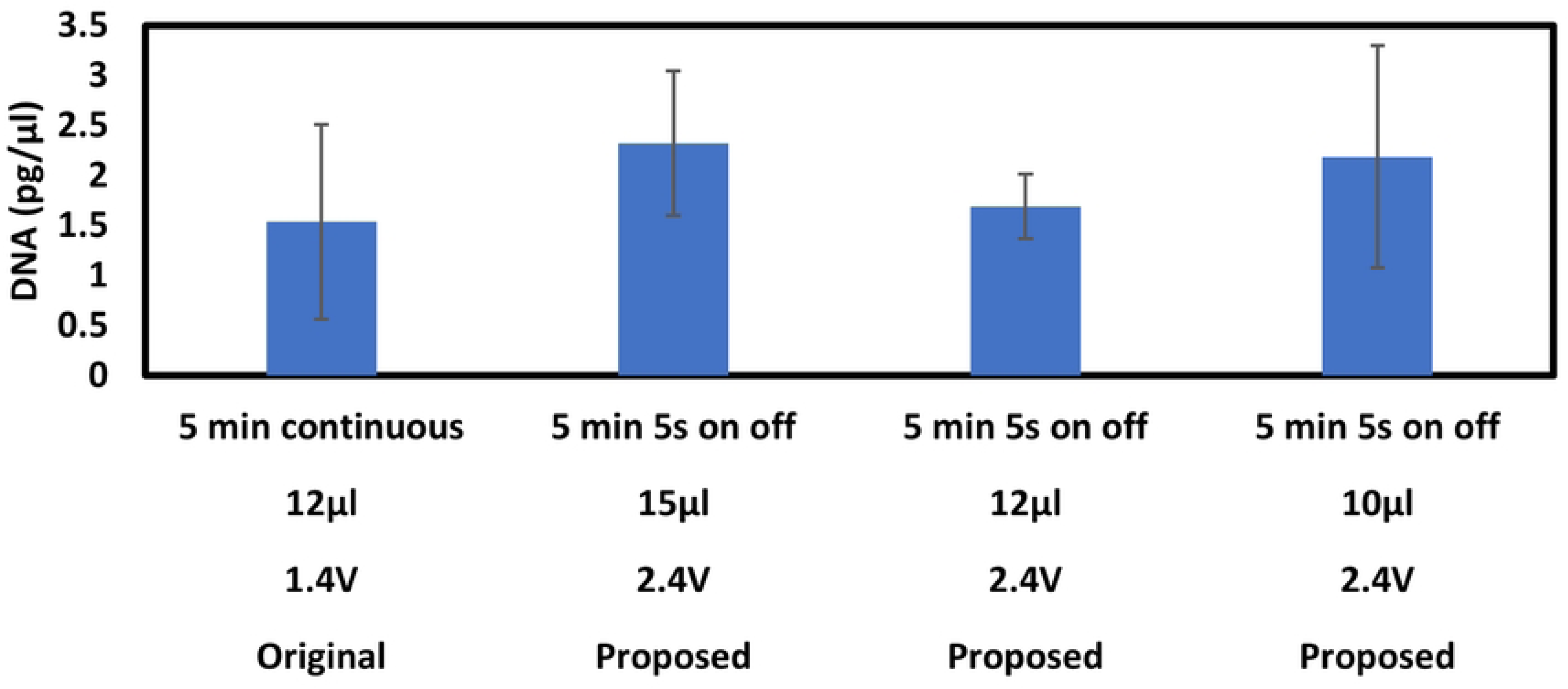
DNA collection for lower volume with a modified chip. (a) Average DNA collection (b) Total DNA collection

#### 3.3.2. Statistical analysis

We repeated the experiments with qPCR three times using both the original and proposed modified chips for different parameters and did not find a statistically significant difference in the repeated tests (P < 0.05). Statistically, when performing a two-sample t-test on the proposed modified chip and the original chip, it was observed (Supplementary Figure 7(a) and 7(b)) that there is no significant difference in average DNA collection between the two chips (p = 0.8). However, when total DNA is considered, we found a significant difference with a p-value < 0.0001. Here, we made no changes to the design other than accommodating for increased volume. This would mean that there is no significant difference in performance due to changes in chip design. However, when considering the total volume, a statistically significant difference is observed, as shown in Supplementary Figure 8(b).

When performing a one-way ANOVA based on operation time, we observed a p-value of <0.0001 for both average and total DNA collection, as presented in Supplementary Figure 8. This indicates that the different time levels: 5 min continuous, 5 min continuous with 5 seconds on/off, 5 min continuous with 15 seconds on/off, and 5 min continuous with 30 seconds on/off have a significant performance difference statistically.

Supplementary Figure 9 presents the one-way ANOVA for the voltage, indicating that the different levels are significant when considering the total DNA collection. For different levels of volume, average DNA collection experiences a statistically significant difference, as indicated by the one-way ANOVA; the results are presented in Supplementary Figure 10.

Supplementary Figure 11 shows the interaction profile for voltage, volume, operation time, and chip design for DNA collection. When volume is plotted against voltage, an opposite trend is observed, where at higher voltages, increasing volume decreases the DNA collection. Whereas for lower voltages, increasing volume improves DNA collection. This demonstrates the effect of frequency, which depends on the applied voltage, on DNA collection as a function of media volume.

When voltage is plotted against volume, DNA collection decreases with increasing voltage at higher volumes, while the voltage effect is weaker at lower volumes. This suggests a modest interaction between voltage and volume.

An interaction plot testing the effect of operation time shows that for 5-minute continuous operation, the DNA collection decreases with an increase in volume. For other instances (such as on/off operation), an increase in volume allows for more efficient DNA collection. Across time, DNA collection increases with increasing voltage for both low and high volumes, with peak collection observed at intermediate operation durations.

The interaction profiles for the original and proposed chip designs follow similar trends across voltage, volume, and time, indicating that chip design does not introduce a meaningful interaction with these parameters.

- The prediction profiler (Supplementary Figure 12) indicates that the proposed chip design, with a voltage of 2.4 V, a volume of 10 µL, and a duration of 5 minutes with 5-second on/off intervals, yields a maximum average DNA concentration of 2.45 pg/µL.

### 3.3.3. Survival data

Figure 10 illustrates the survival rate of embryos 24 hours after ZEG operation for various parameters. The survival rate ranged from 83% to 100% with *n* = 6. No abnormal behavior was observed in the surviving embryos.

**Figure 10.**
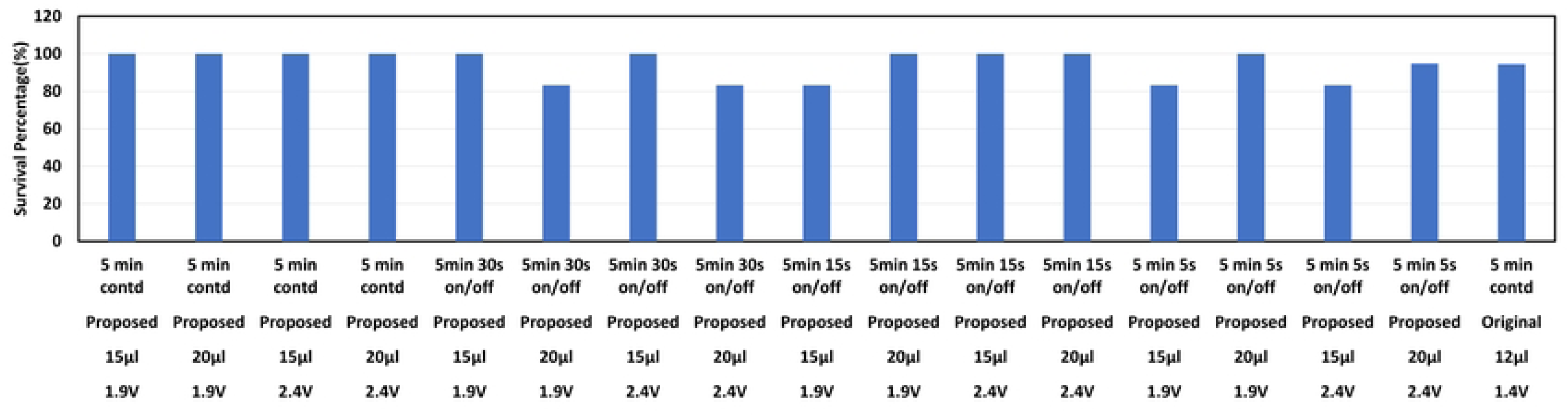
Survival rate of embryos for different values of parameters - voltage, droplet volume, and time of ZEG operation.

## Discussion

In this work, we performed a series of experiments on an automated zebrafish genotyping system (ZEG) focused on the fluid handling chip and its operating parameters in an effort to increase the quantity and concentration of extracted DNA from zebrafish embryos. The experiments were performed on 72-hour post-fertilization (hpf) embryos. The performance was measured using qPCR data to quantify the DNA collection, the amount of fin removal, and the survival of the embryos after using the chip. Changes to the chips that were found to increase the DNA collection included: the surface roughness profile, chip hydrophilicity/hydrophobicity, sample volume, shaker voltage, and the operation time and duty cycle.

The first set of tests were quantified using a fin removal rating to determine the effect of different chip surface roughness profiles on the fin clipping condition of the larva. Two types of roughness profiles were considered with the primary variable being the laser machine’s focusing and defocusing state during roughness generation on the chip. The surface roughness did not cause statistically significant differences in fin removal. However, original profile provided a higher mean value than modified profile, indicating a higher fin removal rating from the ZEG operation. Moreover, these tests with roughness profiles provided information about different parameters that play a vital role and affect the fin removal rating. From the analysis, the sample volume and applied voltage can increase or decrease the fin collection, depending on the level at which they are implemented. The results show that higher sensitivity, or increased fin removal, can be achieved by applying a higher shaker voltage (higher frequency) or using a lower sample volume. The interaction profile indicates that the sample volume and voltage do not interact with each other to a statistically significant extent. However, varying voltage and volume levels can enhance the performance of ZEG operation. After initial testing showed effective fin removal, original roughness profile was considered for further testing. Additionally, evaporation was identified as a serious problem which necessitated changes to the chip design. A proposed modified design, featuring a 3D-printed hydrophilic layer, was found to accommodate a greater volume of water containing the embryo, while still causing the embryos to come into contact with the abrasive surface and was used in the extended testing.

In the fin removal test, we only considered the condition of the fin removal. However, the ultimate performance depends on the quantity of DNA collected for genotyping after the ZEG operation. Fin removal does not entirely represent the amount of DNA collected from the wells; some DNA may become stuck in the rough surface layer and be lost during the collection step. That is why, for the further tests, qPCR was performed to determine the amount of DNA collected as a function of various experimental parameters.

One important parameter that was explored was operation time. From the results, it can be observed that the introduction of on/off (cyclical) operation causes more DNA to be collected compared to continuous operation. Additional experiments on the operating conditions using the modified chip yielded better DNA collection than the original chip with conventional parameters, resulting in a statistically significant improvement. The proposed chip, featuring an improved design and optimized parameters (15 µL volume, 2.4 V, and a 5-minute run with 5 seconds on/off), yielded a DNA collection of 2.68 pg/µL and a total of 38.89 pg. However, the prediction profiler after statistical analysis indicated that 10 µL, 2.4 V, and 5-minute running with 5 seconds on/off as the optimized parameters for the modified chip. Considering the fact that a higher volume is required for our application to solve the evaporation issue, we propose 15 µl, 2.4V, and 5-minute running with 5 seconds on/off as the optimized parameters for the modified chip.

A limitation of the device is that sample loading and collection from the ZEG chip are not automated. Although loading and collection are not time-consuming and only require 2-3 minutes, this can still affect the consistency of the amount collected. The addition of an automated system may contribute to improved consistency.

This ZEG device has only been used for zebrafish embryos. In the future, this technique could be useful with other fish embryos, such as medaka and trout. Moreover, the approach can also be used for larger embryos (>5 days of age) but we have not optimized performance for embryos at that age.

## Acknowledgment

The Author would like to thank the Josh Bonkowsky Lab at the University of Utah School of Medicine, University of Utah and the State of Utah Center of Excellence in Biomedical Microfluidics at the University of Utah for technical support and expertise.

## Author Contribution

Conceptualization: Bruce K. Gale, Raheel Samuel, Christopher Jordan Lambert, Nusrat Tazin

Methodology: Nusrat Tazin, Raheel Samuel

Resources: Bruce K. Gale Software: Bruce K. Gale

Supervision: Bruce K. Gale, Raheel Samuel

Validation: Nusrat Tazin, Christopher Jordan Lambert, Raheel Samuel

Investigation: Nusrat Tazin, Christopher Jordan Lambert, Raheel Samuel

Writing– original: Nusrat Tazin

Writing– review & editing: Nusrat Tazin, Christopher Jordan Lambert, Sabin Nepal, Bruce K. Gale

## Data Availability

The data that supports the findings of this study are available within the article and its supplementary material

